# What do fishermen think? Local perspectives on human–crocodile co-existence in Lake Nasser

**DOI:** 10.64898/2026.04.27.720860

**Authors:** Mohammed A. Ezat, Frank van Langevelde, Marc Naguib

## Abstract

The increasing impact of humans on natural habitats leads to an increase in human–wildlife conflict (HWC), specifically when there is competition for shared resources. In freshwater systems such as Lake Nasser, Egypt, co-occurrence of local fishermen communities with Nile crocodiles (*Crocodylus niloticus*) poses critical challenges for both livelihoods and biodiversity conservation. Understanding local perception of crocodiles by local fishermen is therefore essential for developing effective and socially accepted management responses. We used a structured questionnaire to assess how fishermen perceive and respond to crocodiles across three attitudinal domains: (1) perceived threats, (2) perceived economic benefits, and (3) conservation or co-existence values. Forty-two fishermen were interviewed across multiple khors (side arms of the lake). The responses showed a multifaceted picture: while many local fishermen associated crocodiles with gear damage, reduced catches, and livelihood risks, support for crocodile protection and recognition of their ecological role were also widespread. Interest in crocodile-based livelihood opportunities, such as harvesting or collecting the hatchlings, was generally low, suggesting social, cultural, or legal barriers to such approaches. Fishing experience influenced perceptions, with fishermen encountering crocodiles more frequently reporting decreased catches and greater concern. Cluster analysis further revealed three different respondent groups with different attitudes: conflict-oriented, moderate, and coexistence-oriented. Support for crocodile protection was the strongest predictor of belonging to the pro-co-existence group. Our findings underscore the complexity of human– crocodile co-occurrence in Lake Nasser and, on a wider scale, add to the existing cautions against simplistic mitigations of local HWC. Effective conservation and livelihood interventions will require participatory, context-sensitive approaches that integrate the different perceptions and attitudes of local people.

## 1 Introduction

Humans have an increasing impact on wildlife and their habitat, leading to an increase in human-wildlife conflict (HWC), specifically when there is competition for shared resources and space (Abrahms, 2021; Gaynor and Green, 2026; Blumstein and Fernández-Juricic, 2010). Such conflicts occur across all taxa, from invertebrates (Sayin et al., 2025), fish (Glickman et al., 2025), birds (Noby et al., 2025; Mossad et al., 2023) to larger terrestrial animals such as crop-raiding primates (Hill, 2017) and apex predators that kill livestock and pose dangers to people (Melzheimer et al., 2020; Van Dessel and Snijders, 2023). HWCs are of global concern, yet solutions require understanding of a conflict, taking the biology of the animals as well as the socio-economic situation of the local community into account (Pooley, 2015; Carter and Linnell, 2016; Dickman, 2010; Hull et al., 2023). Understanding the perceptions by the local community indeed is crucial as these underlie people’s actual behaviour and thus affect the acceptance and effectiveness of conservation interventions. Consequently, each type of HWC requires its own understanding, given that the value of sgeneralisations is limited due to the local specificities and complexities of an HWC. Conflicts with apex predators in other regions similarly showed that effects on livelihoods, rather than direct attacks alone, are often the main drivers of negative perceptions among local communities (Akber et al., 2025; Melzheimer et al., 2020).

To better understand the conditions that shape HWC involving free-ranging apex predators, the human-crocodile conflict in Lake Nasser, Egypt, provides an important case. The Nile crocodile (*Crocodylus niloticus*) is a native apex predator whose presence intersects with the livelihoods of many local fishermen operating along the lake’s extensive shoreline (Ezat et al., 2025). While the species is protected under both Egyptian law and international conservation frameworks (CITES, 2026), it is also frequently viewed as a competitor or even a threat by local fishermen, as crocodiles and fishermen co-occur in the same areas (Ezat et al., 2025). This leads to a complex HWC that requires a better understanding of the perceptions of the local fishermen, as shown, for instance, in the conflict of fishermen fishing for crabs and whale entanglements in their nets (Glickman et al., 2025). Thus, in fishermen communities, where direct encounters with crocodiles are potentially hazardous, perceptions are likely to reflect a combination of practical experience, cultural aspects, and economic vulnerability.

Yet, despite the ecological importance of Lake Nasser and the increasing unsystematic reports of human–crocodile conflicts, understanding the perception of local fishermen is missing. Existing studies in the region have largely focused on ecological parameters, not including data on how fishermen themselves experience, interpret, and respond to the presence of crocodiles. For instance, Ezat et al. (2025) showed that the best predictor for crocodile presence is the distance to fishermen’s camps, highlighting the conflict potential through the jointly used space. On the other hand, a survey study was conducted in 2012 to estimate the abundance of the Nile crocodile population in Lake Nasser and to discuss the HWC in broader perspectives, yet without any supporting data and figures (Shirley et al., 2012). Thus, understanding how fishermen in Lake Nasser perceive the presence of crocodiles and whether they consider them as a threat, a resource, or a component of a shared ecosystem is crucial for targeting any conflict mitigation strategies. Studies on snow leopards (*Panthera uncia*) in Pakistan, for instance, found that livestock depredation was the central source of conflict, highlighting how economic vulnerability can strongly shape the attitude toward apex predators (Akber et al., 2025). Without such insights, conservation initiatives and alternative livelihood programs risk being misaligned with local realities. This is particularly relevant in a shared resource system such as fisheries, where livelihoods depend directly on available space and where apex predators cannot be fenced off. In these settings, the perception of wildlife may be jointly shaped by individual and communal social experience, economic vulnerability, and the broader social values (Redpath et al., 2013).

Accordingly, we used a questionnaire for local fishermen to determine fishermen’s perceptions of crocodiles about: (1) perceived threats, (2) potential economic benefits, and (3) conservation or co-existence values. Specifically, we determined whether or not fishermen perceive crocodiles as a direct threat to themselves, what the economic costs and benefits of the presence of crocodiles are, and what attitudes fishermen have towards living together with crocodiles. To do so, we interviewed 42 fishermen in June and July 2023 through a semi-structured questionnaire, using a combination of structured Likert-scale items and open-ended responses. The sample included participants from three fishing associations: Nubian Cooperative Society (n = 21), Fishermen Cooperative Society (n = 18), and Misr Aswan Company (n = 3). We predicted that fishermen would (to a certain degree) see the value of crocodiles, yet that they would also consider crocodiles as a threat to their work. Along this line, we predicted that those who had had negative experiences with crocodiles would see crocodiles as more problematic.

## 2 Methods

We conducted the study in June and July 2023 at Lake Nasser, located in southern Egypt. Lake Nasser is one of the world’s largest man-made reservoirs, extending over 500 km in length and encompassing more than 5,000 km^2^ of surface area, created by the construction of the Aswan High Dam in the 1960s. The lake’s shoreline supports scattered fishing communities operating under cooperative structures regulated by the High Dam Lake Development Authority (HDLDA) (Ezat et al., 2025; Shirley et al., 2012).

### 2.1 Sampling of local sites and fishermen

A total of 42 fishermen were interviewed in multiple camps (N=35) and active fishing zones, including locally more widely known side arms or inlets such as Khor El-Salam, Khor El-Matar, and Khor Abu Ghandour, where crocodile presence and fishing activity overlap. These locations were selected to ensure (1) representation from different fishermen associations with documented crocodile activity (Ezat et al., 2025), (2) geographic diversity along the lake’s western and eastern shorelines, and (3) inclusion of both remote and semi-accessible fishing areas. This spatially explicit sampling strategy thus captured variation in fishermen’s exposure to crocodiles and the surrounding environmental conditions.

We used a non-random sampling strategy to ensure that fishermen were selected who had active and direct exposure to the lake and its wildlife. Site selection was made considering both logistical feasibility and ecological relevance, targeting areas where we expected crocodile encounters to be frequent due to informal reports from local communities.

### 2.2 Data collection

The questionnaire was made in Arabic, pre-tested for clarity, and refined based on feedback from local contacts to ensure cultural and contextual appropriateness. It included several structured demographic and livelihood questions, along with 15 attitudinal statements assessed using a 5-point Likert scale ranging from *strongly disagree* to *strongly agree*. Likert scaling is a standard approach in social research for quantifying attitudes in a scaled way (Jamieson, 2004; Batterton and Hale, 2017; Khadka, 2015).

We collected the data over a six-day period by a field team including the first author and trained local assistants. All interviews were conducted using a face-to-face format, in accordance with best practices for community-based conservation research (Young et al., 2018). We informed the participants about the academic nature of the research, and participation was voluntary. We did not collect any personal identifiers and recorded responses manually using printed questionnaires. Field notes were taken to capture contextual details such as gear damage, presence of crocodile nests, or perceived safety concerns. The questionnaire included structured qualitative demographic and livelihood questions, along with 15 attitudinal statements along a Likert scale. The questionnaire covered qualitative questions concerning: (1) demographics: age, education, household size, (2) fishing profile: gear type, experience, seasonality, cooperative affiliation, (3) crocodile exposure: sightings, incidents, and avoidance strategies, and (4) perception statements. The 15 additional attitudinal statements were categorised into three analytical themes to facilitate structured analysis (Table 1). Responses for these 15 attitudinal statements were recoded numerically (1 = strongly disagree to 5 = strongly agree) and scaled using z-score normalisation to ensure comparability across items. Responses were grouped across three domains: (1) Perceived threats from crocodiles, (2) Perceived economic costs and benefits, and (3) Support for conservation and co-existence (Table 1).

**Table 1.** Thematic classification of 15 Likert-scale statements used to assess fishermen’s attitudes toward crocodiles in Lake Nasser.

| No. | Theme | Statement |
| --- | --- | --- |
| 1 | Threat perception | Crocodiles are a threat to my income |
| 2 | Threat perception | Crocodiles are a threat to me |
| 3 | Threat perception | Fishing in crocodile areas is dangerous |
| 4 | Threat perception | Crocodiles are responsible for reduced fish catch |
| 5 | Threat perception | Crocodiles damage my gear |
| 6 | Economic benefit perception | Harvesting crocodiles provides income |
| 7 | Economic benefit perception | Harvesting crocodile eggs provides income |
| 8 | Economic benefit perception | Harvesting crocodiles would benefit the community |
| 9 | Economic benefit perception | Harvesting crocodile eggs would benefit lake fishermen |
| 10 | Economic benefit perception | Raising crocodiles provides income |
| 11 | Economic benefit perception | Harvesting crocodiles provides more income than fishing |
| 12 | Economic benefit perception | Crocodile-related activities provide continuous income |
| 13 | Conservation / co-existence | Crocodiles and fishermen can live peacefully together |
| 14 | Conservation / co-existence | Crocodiles are part of the natural equilibrium of the lake |
| 15 | Conservation / co-existence | Crocodiles should be protected |

### 2.3 Data Analysis

We digitised all questionnaire responses, calculated descriptive statistics for all Likert-scale items using R (version 2025.09.0), and then summarised responses across three thematic domains (Batterton and Hale, 2017; Jamieson, 2004; Khadka, 2015). To determine how the attitudes of fishermen towards crocodiles were predicted by fishermen responses within the three themes, we compiled patterns in fishermen’s attitudes toward crocodiles using a principal component analysis (unrotated PCA) on five representative indicators selected from across these three domians (Jolliffe, 2002) (Table 2). PCA iswidely used methods for reducing correlated attitudinal variables into principal dimensions of variation (Lever et al., 2017). Following the PCA, we applied a K-means clustering algorithm on the scaled five attitudinal Likert-scaled questions (Table 2) to segment fishermen into groups with similar attitudinal profiles (Jain, 2010). We used the so-called Elbow method to identify the optimal number of clusters (k) by calculating the within-cluster sum of squares (WCSS) across alternative cluster solutions and locating the point at which additional clusters produced diminishing improvements in model fit (Syakur et al., 2018). The elbow point occurred at k = 3, indicating that fishermen’s attitudes were best grouped into three distinct clusters (Figure 4). We then used the scores on the first two principal components (Table 2) to characterize the resulting clusters.

**Table 2.**
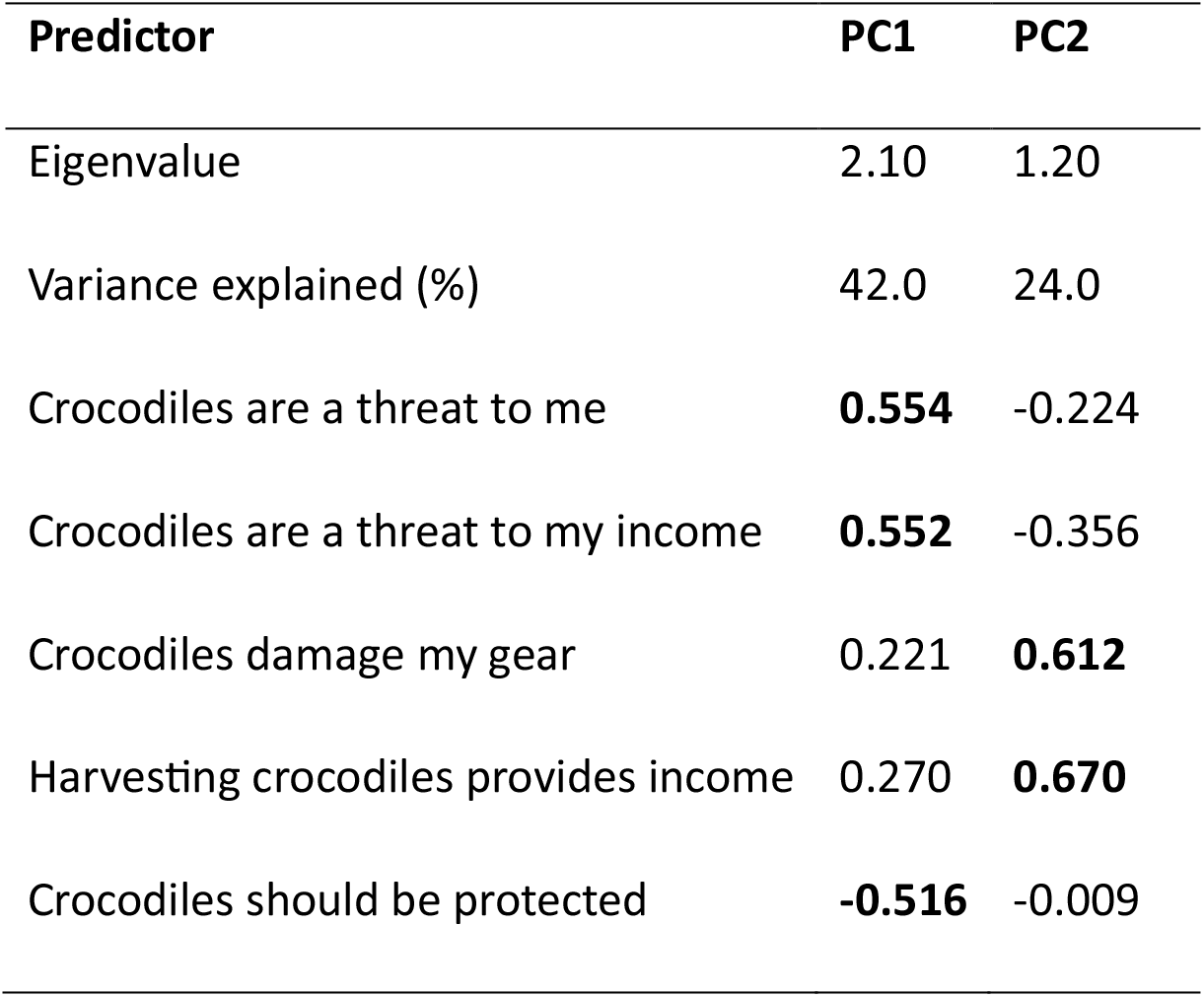
Principal component loadings, eigenvalues, and variance explained for the five Likert-scale statements included in the PCA. 1=strongly disagree, 5= strongly agree) answers to the five questions used in the PCA.

Finally, we determined the individual-level support for crocodile co-existence, based on agreement with the statement: *“*Crocodiles and humans can live together*”* in relation to the three attitudinal themes of this study. The response to the above statement was converted into a binary variable (1 = agree/strongly agree, 0 = otherwise) and modelled as the dependent variable in a binary logistic regression (glm). As independent variables, we selected the Likert scale results from representative questions from the three attitudinal domains to test each research question: (1) threat perception: “Crocodiles are a threat to me” and “Crocodiles are a threat to my income”, (2) economic impacts and livelihoods considerations : “Crocodiles damage my gear” and “Harvesting / killing crocodiles provide income” and (3) conservation attitude: “Crocodiles should be protected”. The analysis for the economic costs incorporated items related to perceived livelihood opportunities (e.g., harvesting or killing crocodiles) as well as reported economic challenges (e.g., damage to fishing gear or fish catch).

## 3 Results

### 3.1 Threat perception

With respect to the threat perception, 43% of the respondents stated that crocodiles pose a threat to their daily income. Similarly, 40% agreed that crocodiles are responsible for reduced fish catch. Regarding personal safety, 38% of the respondents stated that crocodiles are a direct threat to them. Responses to the statement “Fishing in areas where crocodiles are present is dangerous” were mixed with 43% in agreement, 7% neutral, and 43% in disagreement. Finally, a large majority of respondents (80%) agreed that crocodiles damage their fishing gear, while 20 % disagree. These results indicate a diverse range of perceptions regarding both physical and livelihood-related risks, however, many respondents (62%) did not consider crocodiles to be a direct personal threat Finally, gear damage emerged as the most strongly endorsed threat perception item, with 80% of respondents agreeing that crocodiles damage their fishing gear.(Figure 1).

**Figure 1:**
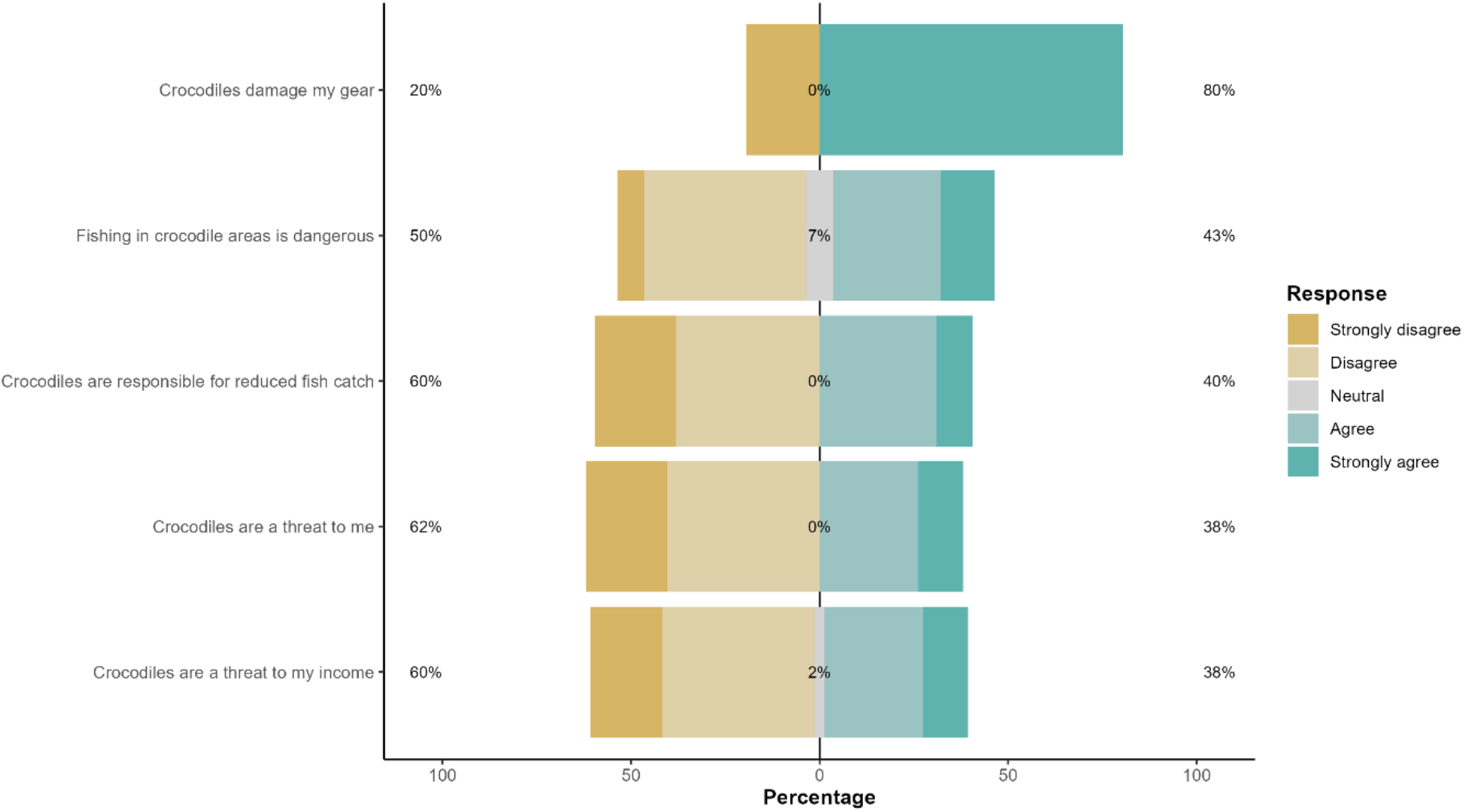
Distribution of responses (%) to five Likert-scale statements related to perceived threats from crocodiles (n = 42).

### 3.2 Perceived economic benefits and costs

Most fishermen did not perceive strong personal economic benefits from illegally harvesting wild crocodiles, collecting eggs/ hatchlings, or engaging in crocodiles related trade. However, perceived benefits for the wider community were higher than perceived benefits for individuals. The majority of the respondents disagreed or strongly disagreed that harvesting wild crocodiles (57%) or collecting their eggs (60%) would be a useful supplementary income source (Figure 2). Similarly, 64% disagreed or strongly disagreed that crocodile use could provide a continuous source of income, while only 10% agreed with this proposition. When asked whether wild crocodile harvesting could generate more income than fishing, 67% isagreed, and only 17% agreed. Perceptions of community-level benefits the local community were more mixed: 29% agreed that wild crocodile harvesting would benefit the local community, while 43% disagreed. A similar pattern was observed for the idea of farming crocodiles for commercial purposes, with 27% in agreement and 38% in disagreement. Neutral responses were common across these items.

**Figure 2.**
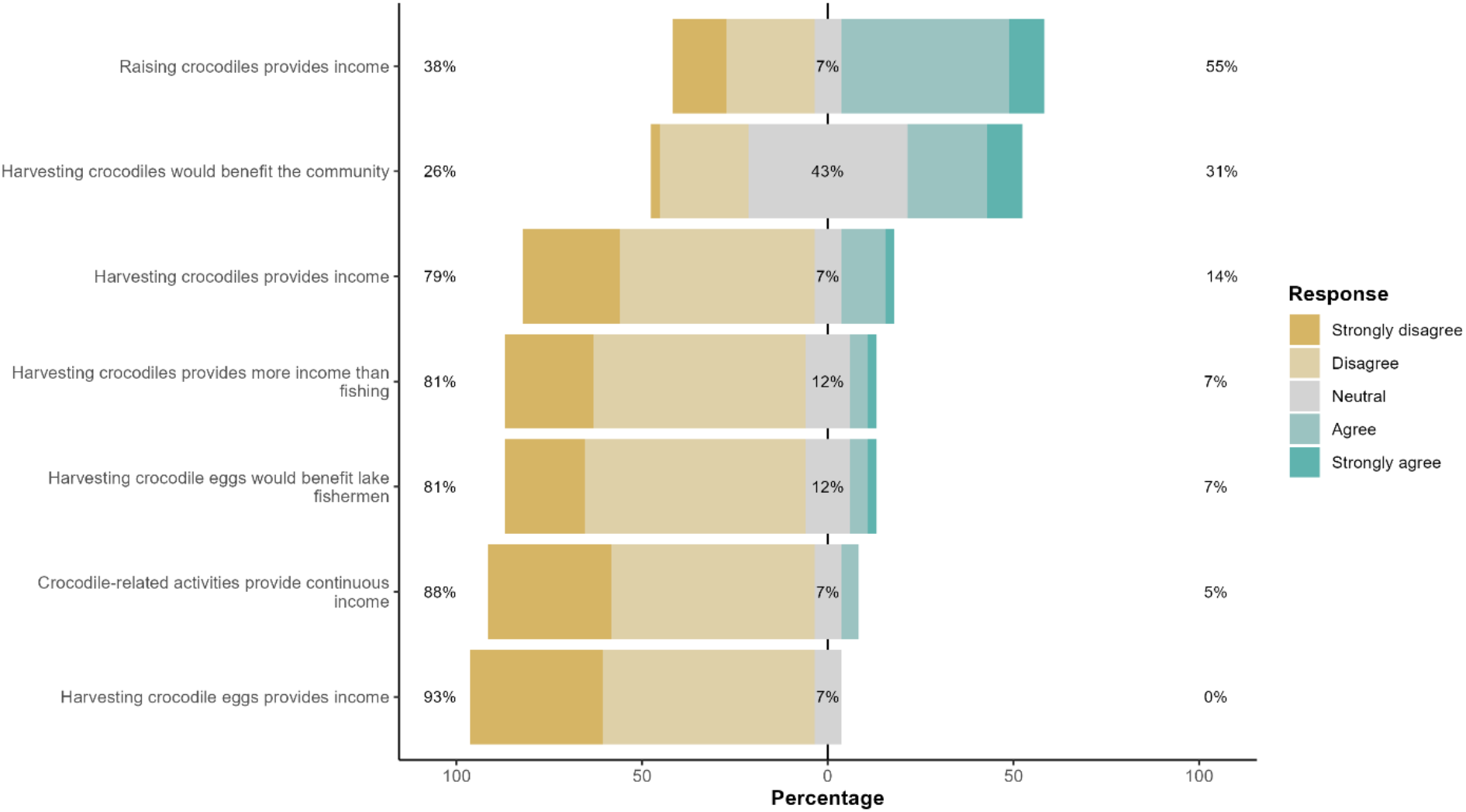
Distribution of responses (%) to seven Likert-scale statements related to perceived economic benefits from crocodile use (n = 42).

Fishermen reported several economic costs associated with crocodile presence. Among respondents who reported frequent crocodile encounters (i.e., daily or weekly), a large proportion indicated that their weekly fish catch had decreased compared to the previous year. By contrast, fishermen who rarely or never encountered crocodiles reported stable or improved catch levels. Some respondents also reported gear loss or net damage attributed to crocodile encounters. In addition, 40% of respondents agreed that crocodiles were responsible for reduced fish catch, while 45% disagreed. Direct engagement with crocodile-related economic activities was limited: fewer than 10% of respondents reported ever capturing or hunting a crocodile, and more than 80% reported no personal involvement in crocodile use (Figure 1, 2).

### 3.3 Co-existence / Conservation perceptions

With respect to the general attitudes, a majority of the respondents (74%) agreed or strongly agreed that crocodiles are part of the natural environment of Lake Nasser (Figure 3). Similarly, 64% agreed that crocodiles should be protected, while 60% of the respondents agreed that crocodiles and fishermen can live peacefully together.

**Figure 3.**
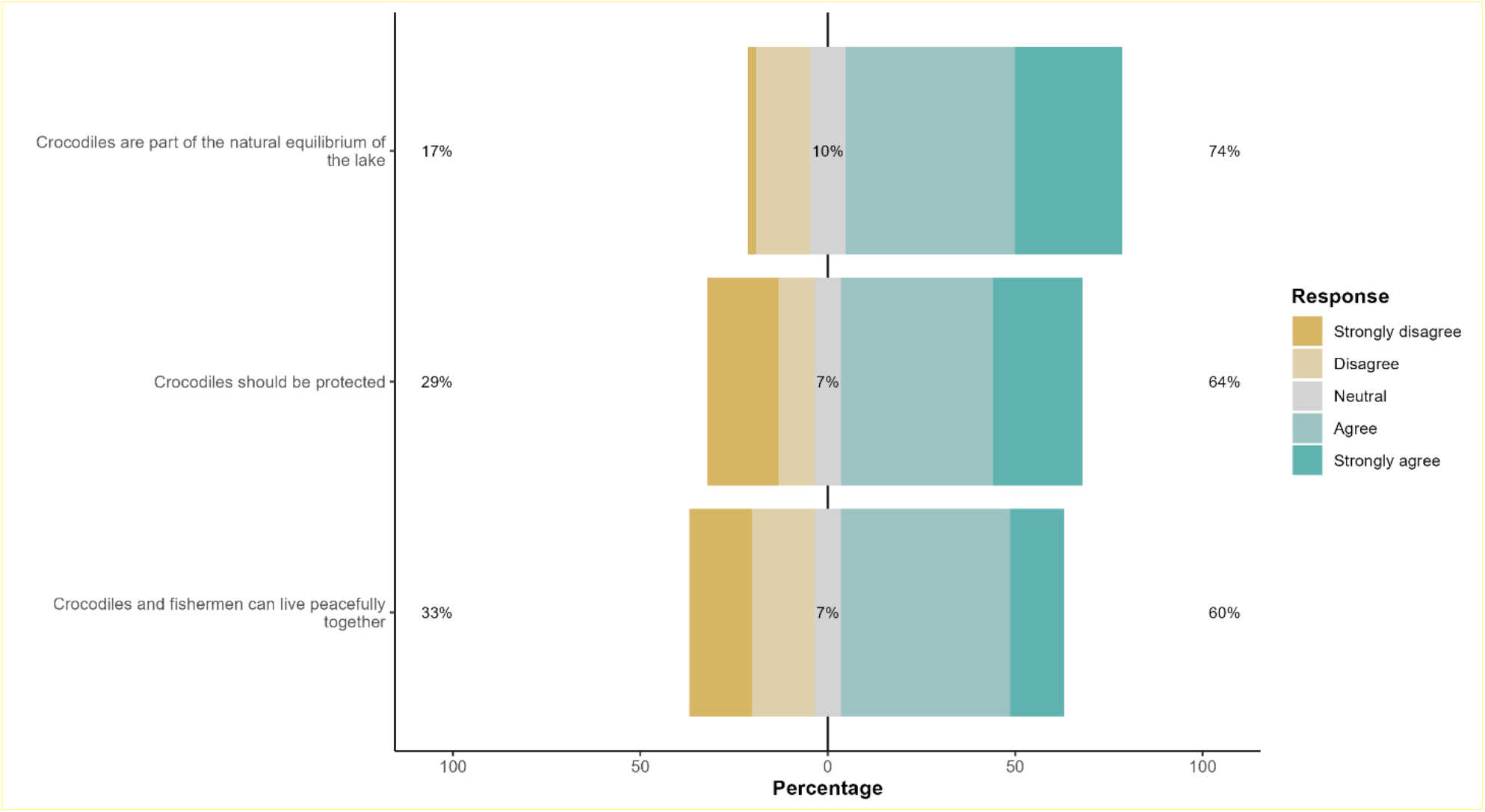
Distribution of responses (%) to three Likert-scale statements related to conservation and co-existence (n = 42).

### 3.4 Fishermen’s attitudes on perceived threats, economic impact, and co-existence

The PCA followed by k means clustering identified three attitudinal groups among fishermen (Figure 4). The conflict-oriented cluster (n = 12) was characterised by stronger perception of crocodiles as a personal and economic threats, greater concerns regarding gear damage, and lower support for co-existence. This group most closely reflects fishermen experiencing pronounced human-crocodile conflict. The pro-conservation cluster (n = 8) showed lower threat perceptions and stronger support for crocodile protection. Respondents in this group generally viewed crocodiles as part of the lake ecosystem and expressed greater acceptance of co-existence. The moderate cluster (n = 21) occupied an intermediate position between the two group. Respondents in this cluster showed more balanced attitude overall, combining relatively low general threat perceptions and support for co-existence while still being concerned regarding practical and economic impacts. At the same time, they were intermediate with respect to support crocodile conservation, suggesting overall a more pragmatic and balanced view of the HWC.

**Figure 4.**
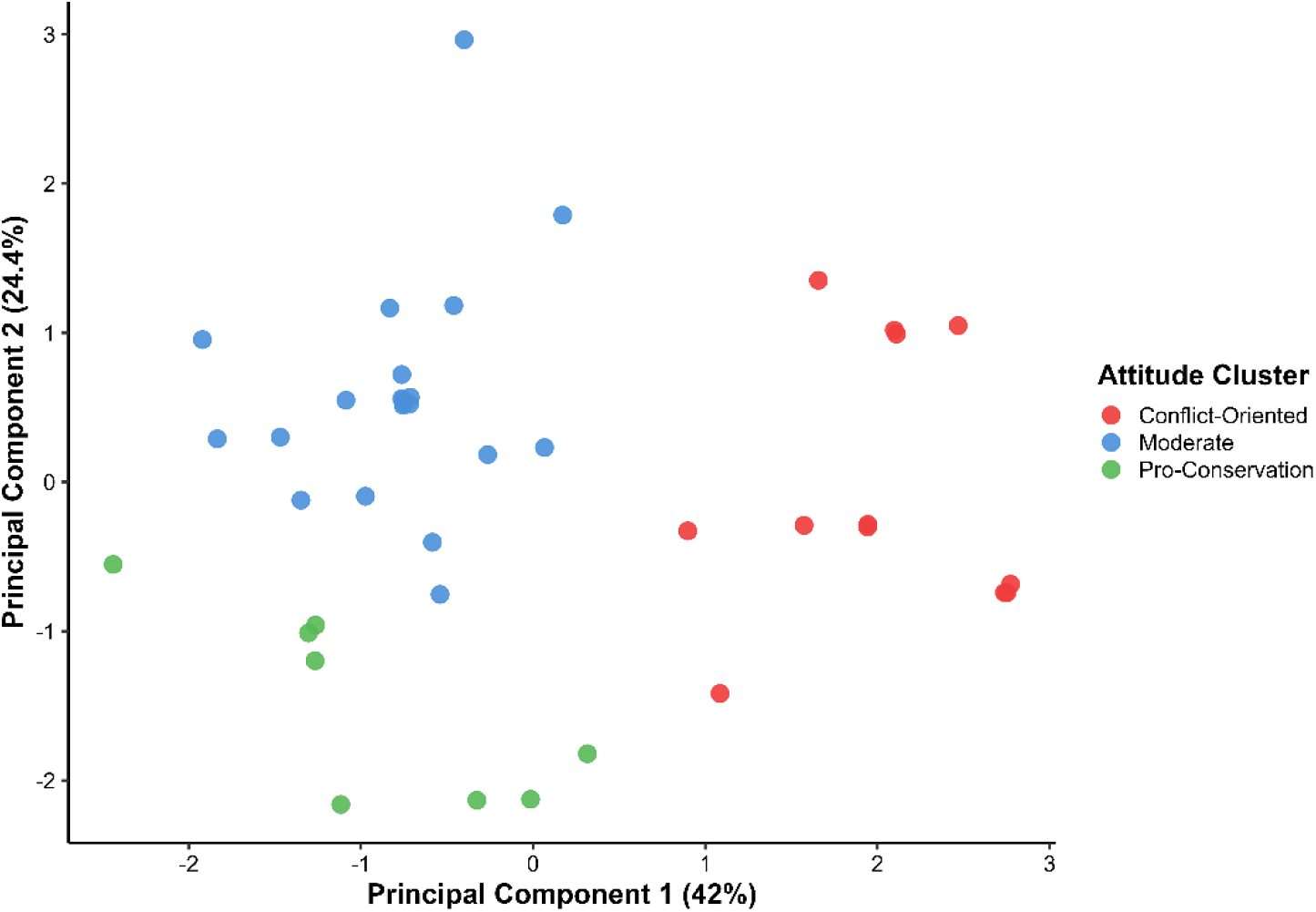
PCA plot showing three clusters of fishermen’s crocodile-related attitudes identified by K-means clustering (k = 3) using scores from the first two principal components (see Table 2). Higher PC1 scores reflect stronger perceived threat and lower support for protection, whereas higher PC2 scores reflect stronger perception of gear damage and economic benefits from crocodile harvesting.

The binomial generalised linear model showed that support for crocodile protection was the only significant predictor of membership in the pro-coexistence cluster (Table 3). Perceived personal threat, income loss, gear damage, and perceived economic benefits of harvesting crocodiles were not significant predictors (Table 3). These results align with the broader PCA-based clustering, reinforcing that fishermen’s views on crocodiles are structured by both risk perception and normative beliefs about wildlife.

**Table 3.** Logistic regression predicting membership in the pro-coexistence attitude cluster.

| Predictor | Estimate | Odds Ratio | 95% CI for OR | p-value |
| --- | --- | --- | --- | --- |
| (Intercept) | -0.272 | 0.76 | 0.00 – 118.54 | 0.916 |
| Crocodiles are a threat to me | -0.589 | 0.56 | 0.20 – 1.37 | 0.209 |
| Crocodiles are a threat to my income | -0.957 | 0.38 | 0.11 – 1.17 | 0.103 |
| Crocodiles damage my gear | 0.056 | 1.06 | 0.53 – 2.18 | 0.874 |
| Harvesting crocodiles provides income | -0.787 | 0.46 | 0.13 – 1.40 | 0.188 |
| Crocodiles should be protected | 1.705 | 5.50 | 1.42 – 28.67 | **0.024** |

**Note:** Odds ratios greater than 1 indicate higher odds of belonging to the pro-coexistence cluster, whereas values below 1 indicate lower odds. Significant effects ( $p < 0.05$ ) are shown in bold.

## 4 Discussion

This study provides one of the first systematic assessments of how fishermen in Lake Nasser perceive and experience free ranging Nile crocodiles, with a focus on the social and economic dimensions of the human-wildlife conflict. The results show that fishermen’s views were shaped less by direct encounters with crocodiles but by gear loss, concerns about income reduction, and broader beliefs about co-existence and conservation. The majority of fishermen were positive toward crocodile conservation and co-existence, and more than 40 % of the fishermen also reported strong concerns regarding the economic use of the crocodiles. These findings highlight the complexity of the fishermen’s perspectives across the three themes examined here, the perceived threat, the economic implications as well as the conservation and co-existence values.

Overall, the findings provide novel insights of how fishermen perceive and engage with free ranging Nile crocodiles and these insights add to the wider perceptions of the complexities of HWCs. Given that a substantial proportion of fishermen reported concerns over crocodiles, particularly regarding gear damage, reduced catch, and fear, their worries about negative effects of the crocodiles likely has direct and indirect effects on their livelihood and economic situation. However, these concerns did not automatically translate into opposition to crocodile conservation in Lake Nasser. Many respondents who recognised the risks still expressed that co-existence is possible. A similar pattern had been observed in tiger (*Panthera tigris*) landscapes, where tolerance varied among local people and was not determined by risk only (Inskip et al., 2016). Therefore, the attitudes towards wildlife are not determined by only direct material costs, but are also influenced by social aspects, cultural acceptance and personal values (Redpath et al., 2013). This pattern is consistent with findings from other apex-predator systems, such as the snow leopard, where material losses to local resource users were identified as a major source of conflict and reducing tolerance towards wildlife (Akber et al., 2025). Moreover, the clustering of attitudes revealed a heterogeneity of views: some respondents fell into a conservation-minded group, others into a conflict-oriented group, and a third group expressed more pragmatic and intermediate views, with in part substantial variation within each cluster. These profiles support the wider view that HWC are not defined by uniform factors, but by complex trade-offs shaped by economic dependencies, personal and social experience, as well as conservation views as also shown for other HWC cases (Dickman, 2010; Gaynor and Green, 2026; Hull et al., 2023). These findings taken together suggest that, as in other HWC cases, any intervention, whether focused on co-existence, protection, or sustainable use, needs to be tuned to this sociological diversity among fishermen to guarantee local acceptance and sustainability. Integrating these sociological insights with an in-depth understanding of the behaviour and ecology of the animals (Clemmons and Buchholz, 1997; Berger-Tal and Blumstein, 2024; Redpath et al., 2013; Sutherland, 1998) is also required to develop mitigation strategies.

Our findings that few fishermen reported involvement in hunting, collecting, or selling crocodile products, and most of them disagreed with statements promoting crocodile harvesting or farming as viable income strategies, imply that any killing or chasing of crocodiles is driven by incentives to protect their fishing, rather than that fishermen target crocodiles out of direct economic needs. This reluctance to seek economic benefits from the crocodiles might stem from multiple factors, such as legal ambiguity, cultural unfamiliarity, limited market exposure, risk aversion as well as from an underlying general acceptance of crocodiles being part of the environment (Dickman, 2010; Redpath et al., 2013). Unlike southern African contexts where community-based crocodile farming has delivered both conservation and economic gains (Boesch et al., 2017), Lake Nasser’s communities lack the enabling conditions for such initiatives. These results caution against assuming that sustainable use models, such as farming, are transferable across ecological or socio-cultural contexts. A recurring concern among fishermen was gear-related damage caused by crocodiles, particularly to fishing nets. This aligns with broader HWC studies, where economic and material losses, rather than direct attacks, represent the most prevalent form of antagonism (Pooley et al., 2017).

Despite material concerns, many fishermen voiced strong support for crocodile protection and acknowledged their ecological role. These views suggest the presence of a positive attitude for co-existence, apparently not grounded in economic benefit but in recognising the ecological role of crocodiles or cultural norms. This aligns with findings from other crocodilian landscapes in South Asia and Africa, where tolerance has persisted even amid occasional conflict (Brackhane et al., 2024). At the same time, economic conflicts do not necessarily eliminate the support for conservation. In the Khunjerab National Park, respondents reported livestock losses through snow leopards, yet favoured improved guarding and predator-proof enclosures rather than killing predators (Akber et al., 2025). Similar patterns have been reported in East Africa, where cultural values among Maasai communities have historically discouraged lion (*Panthera leo*) killing despite conflict with livestock (Hazzah et al., 2017).

Our insights may have several implications, specifically for the conflict between humans and free-ranging wildlife. Conflicts between humans and free-ranging apex predators are complex, not linked to direct land ownership and protection and can involve complex societal dynamics, such as for the wolf (*Lupus lupus*) expanding into human dominated landscapes (Van Dessel and Snijders, 2023). First, interventions aimed at mitigating such conflicts or promoting co-existence must be economically and socially differentiated, as the conflict is not evenly distributed across people, nor are attitudes. Evidence from conflicts involving elephants and large carnivores in other regions shows that effective responses should be adapted to local social and ecological conditions (Dickman, 2010). Second, efforts to introduce livelihood alternatives must be preceded by feasibility assessments, legal frameworks, and market-building efforts. Without these, such proposals are unlikely to work in practice. Moreover, a deeper understanding of crocodile behaviour, their movement patterns, and their relationship to fishermen activities (Ezat et al., 2025) will provide a more holistic view of human-crocodile conflict dynamics. In Lake Nasser and similar systems, bridging ecological and social perspectives is essential for building effective and locally legitimate conservation solutions.

## 5 Conclusions

This study offers the first comprehensive assessments of human–crocodile interactions in Lake Nasser, shedding light on how fishermen perceive, experience, and respond to the presence of Nile crocodiles in their fishing areas. Our findings uncover a diverse pattern with differences in opinions about human-crocodile co-existence shaped by perceived risk, adaptive behaviour, and presumably pragmatism. While crocodiles are frequently associated with gear damage, catch reduction, and behavioural avoidance, most fishermen did not advocate lethal control. Instead, the findings reveal a layered form of tolerance, namely one informed by local ecological awareness, cultural familiarity, and the absence of viable alternatives. Even among those fishermen who experience conflict, there remains a broad recognition of crocodiles as part of the lake’s ecosystem. The findings on the perspectives and attitudes of fishermen in relation to crocodiles support the notion that the human side of HWC is complex and involves multiple views that need to be considered in any mitigation measures (Dickman, 2010).

This research thus underscores the need, also in the HWC between fishermen and free-ranging crocodiles, for nuanced, differentiated conservation approaches that consider local perceptions and recognise diversity within the fishermen community. Combining social science methods with behavioural and ecological data on crocodiles in an interdisciplinary framework will be key for navigating the complexities of human–wildlife co-existence in such a large freshwater ecosystem.

## Acknowledgements

We are grateful for the fishermen participating in the study as well as Alaa Ahmed and Shaezly Abdelfrag for helping in conducting the interviews and Lysanne Snijders for commemts on the mansucript. The research was funded by the Eco2 project awarded to MN by the INREF funds of Wageningen University.

